# Single-shot Echo Planar Time-resolved Imaging for multi-echo functional MRI and distortion-free diffusion imaging

**DOI:** 10.1101/2024.01.24.577002

**Authors:** Zijing Dong, Lawrence L. Wald, Jonathan R. Polimeni, Fuyixue Wang

**Affiliations:** Athinoula A. Martinos Center for Biomedical Imaging, Massachusetts General Hospital, Charlestown, Massachusetts, USA; Department of Radiology, Harvard Medical School, Boston, Massachusetts, USA; Harvard-MIT Health Sciences and Technology, MIT, Cambridge, Massachusetts, USA

**Author notes:** **Correspondence to**: Fuyixue Wang, Ph.D. Athinoula A. Martinos Center for Biomedical Imaging Department of Radiology, Massachusetts General Hospital 149 Thirteenth St., Suite 2301 Charlestown, MA, 02129, USA. **Funding Information:**. This work was supported by the NIH NINDS *BRAIN Initiative* (U24-NS129893), by the NIH NIA (K99-AG083056), and NIBIB (P41-EB030006 and R01-EB019437), and by the MGH/HST Athinoula A. Martinos Center for Biomedical Imaging, and was made possible by the resources provided by NIH Shared Instrumentation Grant S10-OD023637.

**Keywords:** EPTI, single shot, fast imaging, dynamic imaging, distortion-free, multi-echo fMRI, diffusion MRI

## Abstract

**Purpose:** To develop EPTI, a multi-shot distortion-free multi-echo imaging technique, into a single-shot acquisition to achieve improved robustness to motion and physiological noise, increased temporal resolution, and high SNR efficiency for dynamic imaging applications.

**Methods:** A new spatiotemporal encoding was developed to achieve single-shot EPTI by enhancing spatiotemporal correlation in *k-t* space. The proposed single-shot encoding improves reconstruction conditioning and sampling efficiency, with additional optimization under various accelerations to achieve optimized performance. To achieve high SNR efficiency, continuous readout with minimized deadtime was employed that begins immediately after excitation and extends for an SNR-optimized length. Moreover, *k-t* partial Fourier and simultaneous multi-slice acquisition were integrated to further accelerate the acquisition and achieve high spatial and temporal resolution.

**Results:** We demonstrated that ss-EPTI achieves higher tSNR efficiency than multi-shot EPTI, and provides distortion-free imaging with densely-sampled multi-echo images at resolutions ∼1.25–3 mm at 3T and 7T— with high SNR efficiency and with *comparable* temporal resolutions to ss-EPI. The ability of ss-EPTI to eliminate dynamic distortions common in EPI also further improves temporal stability. For fMRI, ss-EPTI also provides early-TE images (e.g., 2.9ms) to recover signal-intensity and functional-sensitivity dropout in challenging regions. The multi-echo images provide TE-dependent information about functional fluctuations, successfully distinguishing noise-components from BOLD signals and further improving tSNR. For diffusion MRI, ss-EPTI provides high-quality distortion-free diffusion images and multi-echo diffusion metrics.

**Conclusion:** ss-EPTI provides distortion-free imaging with high image quality, rich multi-echo information, and enhanced efficiency within comparable temporal resolution to ss-EPI, offering a robust and efficient acquisition for dynamic imaging.

## Introduction

Capable of generating a 2D image in a single shot, Echo-Planar Imaging (EPI) (1) has been the standard sequence for various key neuroimaging applications, including functional, diffusion, and perfusion MRI. The development of EPI-based multi-echo (2–6) and multi-contrast (7–10) sequences has further expanded its utility, providing additional information about brain function and structure. For instance, multi-echo imaging has been demonstrated to be useful for enhancing functional sensitivity (11–14) and removing undesirable physiological noise in fMRI (15–19). For diffusion imaging, multi-echo images are valuable for multi-compartment modeling and diffusion-relaxometry analyses, offering more specific and comprehensive measurements of tissue microstructure (20–23). The integration of gradient and spin multi-echo sequence has also been used to achieve fast T_2_ and T * mapping and measure vascular properties in perfusion imaging (8–10,24,25).

However, EPI or EPI-based techniques suffer from several limitations that compromise both image quality and information content. These limitations include geometric distortion along the phase-encoding (PE) direction due to *B*_0_ inhomogeneity (26,27), which compromises anatomical integrity and leads to resolution loss. This geometric distortion can also dynamically vary over time due to *B*_0_ field variations caused by changes in head position (27), field drift (28), or physiological dynamics (e.g., the respiratory cycle) (29,30), thereby compromising the temporal stability of the image time series. The T_2_/T * decay across the EPI readout also leads to additional limitations, including spatial blurring, signal dropout, and unwanted contaminations into the image contrast at the desired TE (31,32). Finally, the extended image readout time limits the acquisition of multi-contrast/multi-echo images before the signal decays away. Considerable efforts have been made to alleviate these limitations. The most common acquisition methods include parallel imaging (33–35) and multi-shot EPI techniques (36–43), both of which reduce the effective echo spacing and readout length, therefore reducing distortion/blurring and increasing the number of echoes that can be acquired. However, these approaches cannot fully address these challenges, and their ability to mitigate these problems often comes at a cost of high noise amplification and aliasing artifacts, or leads to trade-offs in temporal resolution and increased vulnerability to physiological noise, especially when using large acceleration or multi-shot factors. For distortion mitigation, methods that combine reversed phase-encoding acquisition with distortion correction have also been developed (44–46).

Echo Planar Time-resolved Imaging (EPTI) (47) was first introduced as a multi-shot technique to address the inherent limitations of conventional EPI. While parallel imaging and multi-shot EPI alleviate these problems by reducing the phase and magnitude change across a shorter readout, EPTI takes a different approach. It recognizes the strong spatiotemporal correlation of the data within the readout and efficiently encodes in a spatiotemporal domain, which allows it to replace the conventional EPI image formation and resolves multi-echo images free from artifacts associated with the accumulated imperfections. This turns a liability—the evolution of the MR signal with time, which yields distortion/blurring—into an asset, providing benefits and unique features—distortion- and blurring-free, multi-echo images with a wide range of TEs, and high readout efficiency. The original multi-shot EPTI (ms-EPTI) mostly targets anatomical imaging and high spatial resolution applications. For example, it has been demonstrated to provide high efficiency for fast multi-contrast imaging (47,48), multi-parametric quantitative MRI (49,50), high-resolution functional MRI (51), high-resolution distortion-free diffusion imaging (52–54), and myelin water imaging (55).

While multi-shot EPTI is highly suitable for high spatial resolution imaging, many dynamic imaging applications, such as functional, diffusion, and perfusion MRI, require a single-shot acquisition for high robustness to motion and physiological noise and/or high temporal resolution. Therefore, this work aims to exploit EPTI in the single-shot regime, and develop a robust and practical single-shot EPTI (ss-EPTI) acquisition for dynamic imaging applications. Our goal is a new acquisition tool that overcomes the limitations of ss-EPI, while providing improved image quality, higher efficiency, and multi-contrast capability.

To achieve single-shot EPTI, our approach, first described in the abstract form (56,57), introduces a new spatiotemporal encoding pattern to provide enhanced spatiotemporal correlation in the *k_y_*-*t* space compared to the conventional EPTI. It avoids large gradient blips and minimizes echo-spacing (e.g., ∼40% shorter at 2-mm resolution) under the single-shot regime, therefore achieving improved reconstruction conditioning and sampling efficiency. We evaluate the proposed single-shot encoding at different acceleration factors and spatial resolutions to identify an optimized encoding strategy for high reconstruction accuracy. In addition, *k-t* partial Fourier acquisition is integrated with ss-EPTI to further accelerate the acquisition and achieve higher spatial resolution. Simultaneous multi-slice (SMS) (58,59) acquisition is also employed in ss-EPTI to further improve temporal resolution.

Leveraging these developments, we show that single-shot EPTI now provides images free from distortions with densely-sampled multi-echo images at isotropic resolutions ranging from ∼1.25–3 mm at 3T and 7T—with high tSNR efficiency and *comparable* temporal resolution to ss-EPI. We demonstrate that single-shot EPTI provides higher temporal SNR (tSNR) than ms-EPTI owing to its improved robustness against motion/physiological noise. Compared to ss-EPI, ss-EPTI also achieves high tSNR efficiency by using continuous readout to fill in any dead time in the sequence. Since a longer readout no longer comes at the cost of severe distortion/blurring as in ss-EPI, the flexible EPTI readout can begin immediately after RF excitation (e.g., an early TE of 2.9 ms) and extend for a duration chosen to achieve optimal SNR efficiency. Furthermore, the ability of ss-EPTI to provide time-series free from dynamic distortions present in EPI can further improve temporal stability, particularly during acquisitions with large field changes due to physiological dynamics such as breathing.

While ss-EPTI can serve as a general readout technique for various imaging applications, this article focuses primarily on demonstrating its efficacy in the context of multi-echo fMRI and distortion-free diffusion imaging. For multi-echo fMRI, we show that ss-EPTI’s early TEs can be used to recover signal intensity and functional sensitivity dropout in challenging brain regions, whose signal can be irrevocably lost in ss-EPI. We also demonstrate the feasibility of combining ss-EPTI with Multi-Echo Independent Component Analysis (ME-ICA) (17,60,61) denoising to distinguish non-BOLD components from BOLD signals by exploiting the TE-dependent information, thereby improving tSNR. In diffusion MRI, ss-EPTI can minimize spin-echo TEs by sampling immediately after diffusion encoding to improve SNR efficiency, and provides high-quality diffusion-weighted images free from both geometric distortion and eddy-current dependent distortion, as well as multi-TE diffusion metrics. These results underscore the promise of ss-EPTI for various dynamic imaging applications.

## 2 Theory

### 2.1 Single-shot EPTI acquisition

Conventional EPI generates a single distorted and blurred image after combining *k*-space samples evolving across time with accumulated *B*_0_ phase and signal decay (Fig. 1A left). EPTI (47) samples the *k-t* (TE) space with highly accelerated spatiotemporal encodings (Fig. 1A middle), and exploits the strong data correlation to fully reconstruct the *k-t* space, generating >100 distortion/blurring-free multi-echo images across the readout. The original EPTI implementation acquires different *k_y_*segments in a multi-shot manner, with each shot consisting of multiple *k_y_*-line sections (the orange rectangles with dashed lines) to ensure regular sampling to apply *k-t* GRAPPA kernel (47,62), and achieves small gradient blip sizes to reduce echo spacing (ESP) and eddy currents. However, when moving directly to single-shot acquisition, the original EPTI encoding pattern would require large gradient blips to jump through the entire *k-*space from −*k_y_*_,max_ to +*k_y_*_,max_, leading to increased ESP, dead time, and eddy currents. The increased ESP and dead time will not only reduce the readout efficiency but also lower the temporal correlation of the sampled data within the readout, resulting in high noise amplification and image artifacts after reconstruction.

**FIG. 1.**
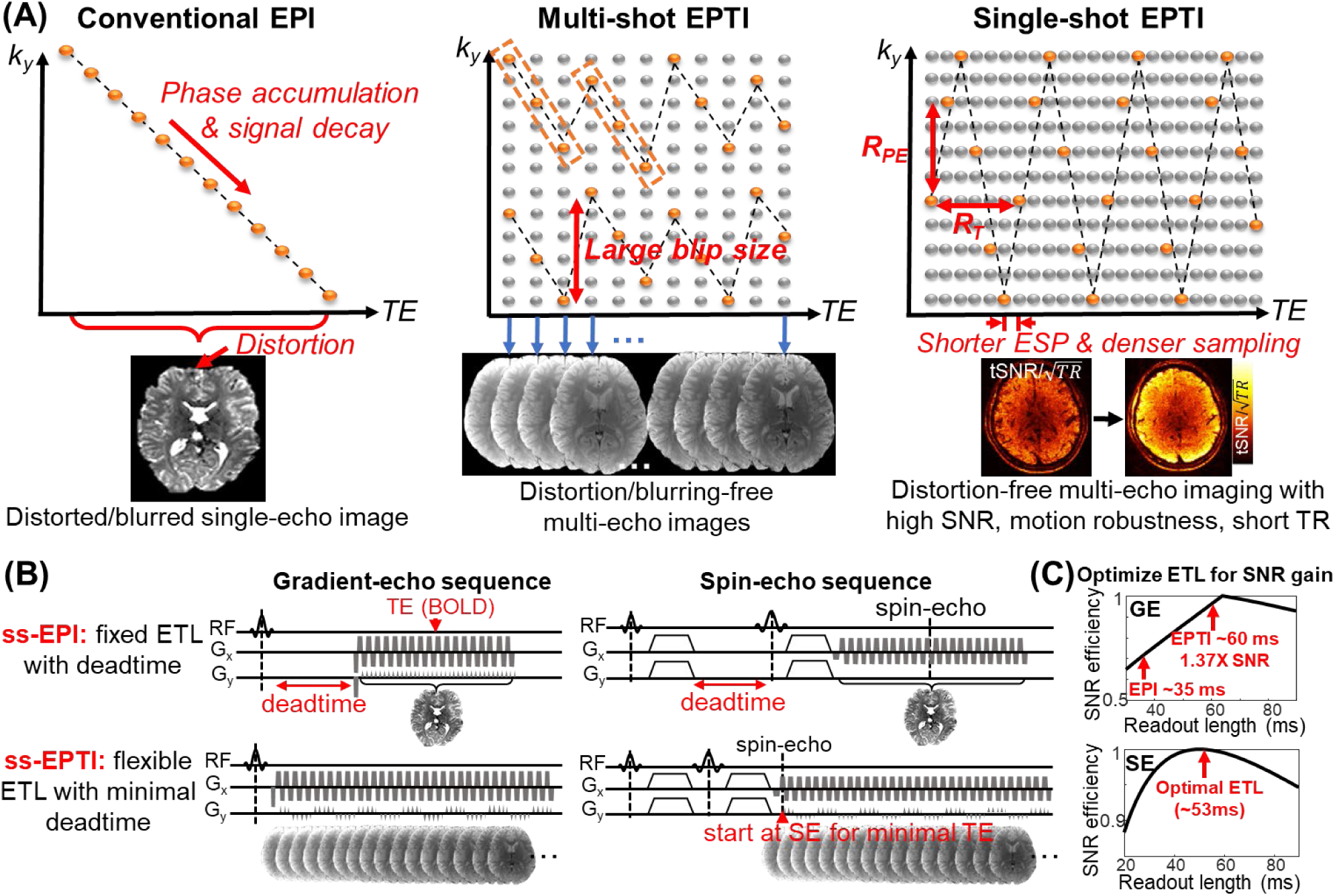
(**A**) Illustration of encoding in conventional EPI, multi-shot EPTI, and single-shot EPTI in *k-t* space. Conventional EPI acquires different phase-encoding (*k_y_*) lines at different time points, and produces a single distorted and blurred image after combining these PE lines with accumulated phase and signal decay. EPTI efficiently samples the *k_y_*-*t* space with highly accelerated encodings in a few shots, and exploits the spatiotemporal correlation to recover the full *k_y_*-*t* data, resolving >100 distortion-free multi-echo images. Single-shot EPTI was developed to acquire the *k_y_*-*t* data using only one shot. The encoding pattern is designed to enhance spatiotemporal correlation to enable robust single-shot reconstruction. It continuously traverses through the *k-t* space with minimal gradient blips required, reducing dead time and ESP compared to conventional EPTI encoding in the single shot regime, resulting in improved reconstruction conditioning and sampling efficiency. As a result, distortion-free multi-echo imaging can be achieved with high SNR efficiency, high robustness to motion/physiological noise, and high temporal resolution. (**B**) Illustration of the SNR-efficient continuous readout employed in ss-EPTI in both gradient-echo (top) and spin-echo (bottom) sequences, which fills in dead time usually wasted in conventional EPI and provides high readout efficiency. Unlike EPI, ss-EPTI gains SNR from a longer readout without introducing severe distortion/blurring or being constrained by spatial resolution as seen in conventional EPI. (**C**) Theoretical SNR efficiency (SNR/√TR) as a function of readout length for gradient echo and spin-echo ss-EPTI sequences. The plots show that the readout length can be optimized in ss-EPTI for optimal SNR efficiency. The red arrows indicate typical readout lengths in ss-EPI and ss-EPTI, highlighting the increased SNR efficiency using an optimized length.

#### Single-shot EPTI encoding with enhanced spatiotemporal correlation

To address this issue and achieve robust single-shot EPTI acquisition, we developed a zig-zag trajectory (Fig. 1A right) that continuously traverses through the *k-t* space with minimal gradient blips required, thereby reducing the dead time and achieving shorter echo-spacing. For example, at a 2-mm resolution, this new ss-EPTI encoding strategy can provide ∼40% shorter ESP (from 0.93 ms to 0.55 ms) compared to the previous encoding pattern. This encoding with reduced echo-spacing, on the one hand, enhances spatiotemporal correlation in *k-t* space and reduces g-factor and artifacts in the EPTI reconstruction. On the other hand, the reduced dead time leads to denser sampling and increases the sampling efficiency. The combination of the g-factor reduction and denser sampling leads to higher SNR efficiency. In addition, complementary *k_y_* sampling across time is employed, in which different *k_y_* points are sampled during each pass through *k*-*t* space. The acceleration factor along *k_y_* (*R*_PE_, defined as the distance along *k_y_* between the consecutive samples) and along time (*R*_T_, temporal distance between *k_y_* samples) are interdependent and synergistically determine the spatiotemporal correlation of the single-shot *k-t* data. A balanced selection of *R*_PE_ and *R*_T_ can further improve reconstruction performance (Fig. 2B). Specifically, a ‘spatial-favored’ configuration with a small spatial undersampling factor *R*_PE_ and a large temporal distance factor *R*_T_ results in weak temporal correlation, hindering the ability to take advantage of the complementary *k_y_* sampling across *t*. Conversely, a ‘temporal-favored’ setup with a high *R*_PE_ and a small *R*_T_ challenges the capability of utilizing the spatially-varying coil sensitivity profiles to estimate missing samples even with the temporally complementary sampling. Based on these considerations, an optimized encoding strategy can be identified that balances the spatial and temporal acceleration to best leverage spatiotemporal correction.

**FIG. 2.**
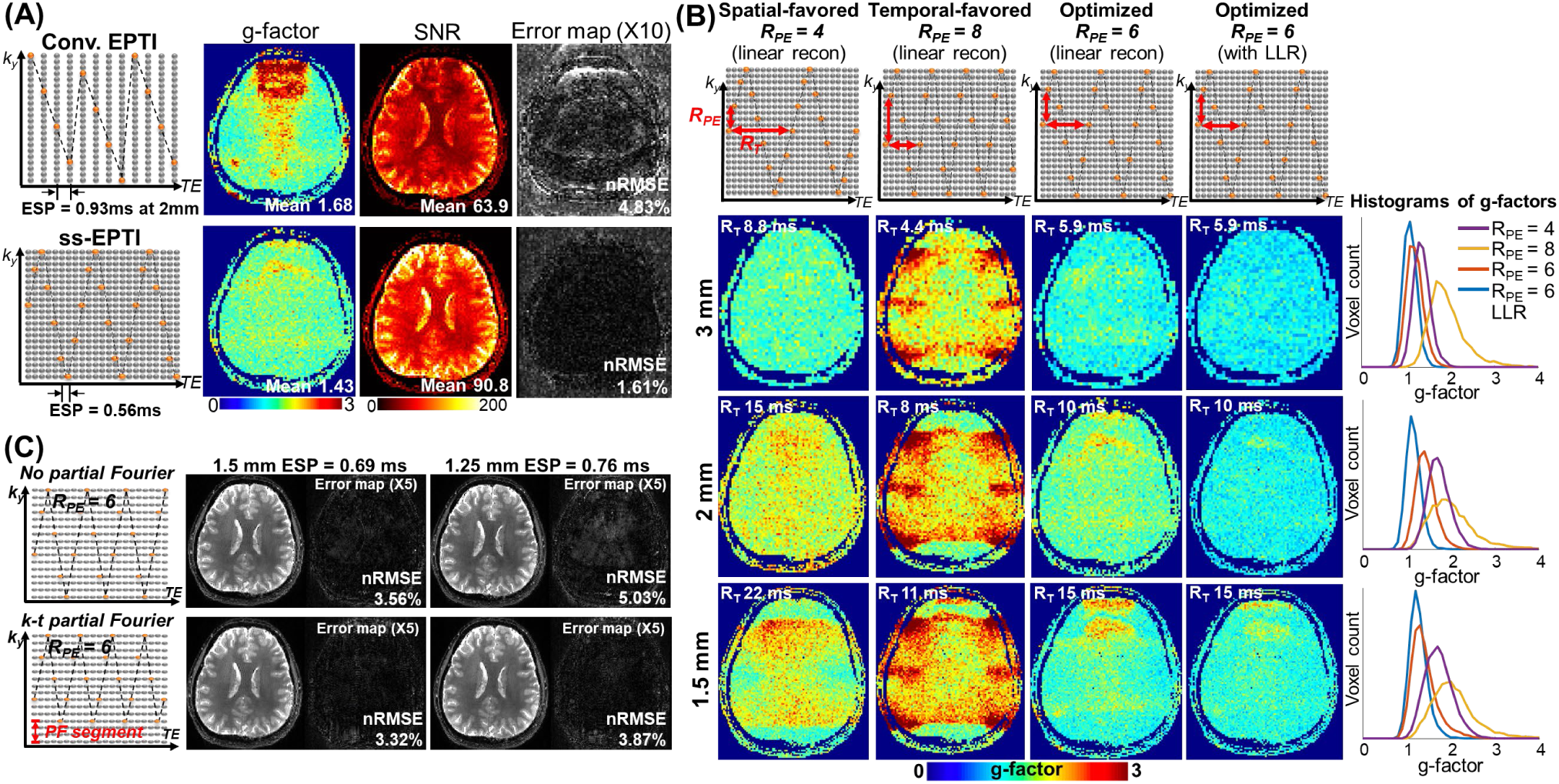
Evaluation and optimization of the proposed encoding through simulation experiments. (**A**) Comparison between the conventional and proposed single-shot encoding using a Monte Carlo simulation at 2-mm resolution. The proposed single-shot encoding reduces the ESP from 0.93 ms to 0.56 ms in comparison to the conventional encoding. Maps of g-factor, SNR, and reconstruction error (scaled by 10×) calculated from the echo-averaged signal are presented for both encodings. (**B**) G-factor maps of the proposed ss-EPTI encoding at different spatial (*R*_PE_) and temporal (*R*_T_) acceleration factors, evaluated at 3-mm, 2-mm, and 1-mm spatial resolutions. Linear subspace reconstruction was used for SNR/g- factor calculation. An instance of the reconstruction with LLR regularization is also included to show the noise level after applying the LLR constraint, but note that it is specifically valid within the tested noise level and image characteristics and may not be universally applicable due to the non-linear nature of LLR regularization. The last column displays the histogram of the g-factor values for all presented cases. (**C**) Single-shot EPTI encoding without and with *k-t* partial Fourier acquisition, designed to improve reconstruction accuracy at higher spatial resolutions. Corresponding reconstructed images (echo-combined) and error maps (×10) at 1.5 mm (ESP = 0.69 ms) and 1.25 mm (ESP = 0.76 ms) resolution are shown.

#### Continuous single-shot readout for high SNR efficiency

Since a longer readout no longer comes at the cost of severe distortion/blurring nor is it constrained by resolution as in ss-EPI, ss-EPTI can improve readout and SNR efficiency by using a continuous readout to fill in dead time in the acquisition that is usually wasted in conventional EPI (e.g., wait time required to achieve the typical TE value for BOLD-weighted fMRI, or dead time between excitation and refocusing pulse in a diffusion-weighted spin-echo sequence) as shown in Fig. 1B. The EPTI readout begins immediately after RF excitation/refocusing (e.g., a very early TE of 2.9 ms) and extends for a duration chosen to achieve optimal SNR efficiency. As shown in Fig. 1C, using gradient-echo and spin-echo sequences as examples, a longer readout window achieves higher SNR efficiency by reducing the thermal-noise level through noise averaging of more signals. By achieving an optimal readout length that surpasses the typical readout length used in ss-EPI due to distortion/resolution constraints, ss-EPTI can achieve higher SNR efficiency. Beyond the optimized length, SNR efficiency decreases because further increase in readout length lengthens the TR and only samples decayed signal with lower intensity.

#### Comparable temporal resolution to EPI and further combination with SMS

By achieving single-shot acquisition, EPTI can now achieve comparable temporal resolution to single-shot EPI, while providing distortion-free imaging and rich multi-echo information. For example, a single TR timepoint (e.g., ∼3 seconds without SMS) in a gradient-echo ss-EPTI protocol (e.g., 3-mm iso), generates >100 whole brain distortion-free volumes, each with a distinct TE in the range of 3–60 ms spaced at an ESP of 0.5 ms. This is in comparison to EPI with equivalent spatial coverage and resolution that achieves a similar TR of ∼3 s for a single-echo volume. To further reduce the volume TR, ss-EPTI can be combined with simultaneous multi-slice (58,59). Specifically, multiband RF excitation is applied and a *k_y_-k_z_-t* sampling trajectory is employed: a *k_z_* blip is applied after each *k_y_* sampling, with alternating blip polarity similar to the blipped-CAIPI trajectory (59), sampling *k_z_* positions with good sampling coverage in *k_y_-k_z_-t* space.

#### k-t partial Fourier acquisition for acceleration at high spatial resolution

A *k-t* partial Fourier acquisition for EPTI is introduced to further accelerate the acquisition in the *k-t* space and to achieve higher spatial resolution (Fig. 2C). In contrast to conventional EPI, EPTI does not require conventional partial Fourier acquisitions to reduce TE or readout length, because its time-resolving feature already enables multi-echo imaging with a flexible TE range and readout length. Instead, the *k-t* partial Fourier encoding with asymmetric sampling along the PE direction is proposed to achieve higher resolution *without* necessitating an increase in the acceleration factor. In other words, when aiming for a high spatial resolution, the *k-t* partial Fourier encoding can help reduce the acceleration and improve reconstruction conditioning. During the reconstruction, the sampled area is initially reconstructed using subspace reconstruction (described below), and subsequently transformed into *k*-space to execute a POCS partial Fourier reconstruction (63) for all echoes to preserve spatial resolution.

### 2.2 Time-resolved subspace reconstruction for ss-EPTI

A subspace reconstruction (48,64–66) is used to reconstruct the multi-contrast images of ss-EPTI. This subspace approach, previously evaluated in our multi-shot EPTI studies (48), has demonstrated robust performance for highly accelerated data. Compared to the original GRAPPA-based reconstruction (47,62), the subspace approach can reduce the number of unknowns by representing the temporal signal evolution using a set of subspace basis and reconstructing the coefficients of the subspace basis set:

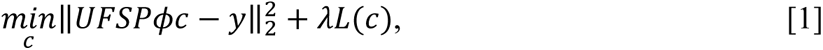

where *ϕ* is the subspace basis, *c* is the coefficient maps of the subspace basis set, *P* is the phase term of the EPTI data including the *B*_0_-inhomogneity and eddy-current induced phase, *S* is the coil sensitivity, *F* represents the Fourier transform, 𝑈 is the undersampling mask in *k-t* space, and *y* is the acquired EPTI data. *L*(*c*) is the locally low-rank (LLR) regularization applied to the coefficient maps, and 𝜆 is its control parameter. Here, when using the LLR constraint, the temporal basis elements are scaled based on their singular value computed during basis generation to balance the amplitude of different coefficient maps, improving the performance of the LLR regularization. For SMS acquisitions, the simultaneously acquired slices are reconstructed by expanding the model to include SMS encoding in both *F* and *U*.

A fast low-resolution *k-t* calibration scan is acquired to estimate the coil sensitivity (*S*) and the phase term (*P*). The calibration scan is fully sampled along time (i.e., along TE), and accelerated along *k*_y_ in the *k*-space periphery to reduce calibration scan time, with a fully-sampled center for autocalibrated GRAPPA (35) reconstruction. Both the *B*_0_ map and the odd-even-echo phase difference map (caused by eddy currents from alternating readout gradient polarities, leads to ghosting artifacts in EPI) are estimated from the calibration data and modeled into the reconstruction. The *B*_0_ map is updated every TR to capture dynamic field changes due to drift or physiological changes (e.g., respiration). This *B*_0_ field change can be estimated through a pre-reconstruction of ss-EPTI data with *B*_0_ phase change modeled in the subspace basis set (52), or simply from the first and third echoes of the standard 3-line navigator (51).

## 3 Methods

Experiments were performed on a Siemens Prisma 3T scanner with a 32-channel head receiver coil (Siemens Healthineers, Erlangen, Germany) and a Siemens Terra 7T scanner with 32-channel Nova coil (Nova Medical, Wilmington, MA). *In-vivo* data were acquired on healthy volunteers with a consented institutionally approved protocol. The BART toolbox (67,68) was used to perform the EPTI reconstruction. We have made the EPTI sequences and reconstruction available (see EPTI dissemination website: https://martinos.org/~fw089/, and MGH C2P website: https://nmr.mgh.harvard.edu/c2p) upon request, including both single-shot EPTI and multi-shot EPTI sequences with example protocols. The reconstruction code is also available for download on Github (https://github.com/FuyixueWang/EPTI).

### 3.1 ss-EPTI encoding optimization

To optimize the spatiotemporal encoding for ss-EPTI, Monte-Carlo simulations were first performed to assess the performance of different encoding strategies in terms of the g-factor, SNR, and reconstruction accuracy. While the g-factor evaluates the noise amplification due to the reconstruction compared to the fully-sampled *k-t* data, SNR also assesses the data sampling efficiency of the readout (e.g., an encoding pattern with a shorter ESP and smaller blips will sample more data in the same readout duration with less inter-sample dead time, resulting in higher SNR). The reconstruction accuracy was measured using normalized root mean square error (nRMSE) calculated between the fully-sampled reference data and the reconstructed images of the undersampled data. EPTI data were simulated using a set of pre-acquired proton density, T_2_*, and *B*_0_ maps. Simulated fully-sampled 32-channel *k-t* space data were generated with added random noise (SNR=40), which was then undersampled with different encoding patterns. The ESP of the simulated data was determined using values realistically achievable at the corresponding spatial resolution and the blip size required by the encoding pattern. The readout duration was held constant at ∼60 ms for all cases to ensure a fair comparison in SNR. The conventional EPTI and ss-EPTI encodings at different spatial resolutions (1.5–3 mm) and spatiotemporal acceleration factors (*R*_PE_ and *R*_T_) were evaluated (for details of encoding parameters, see Fig. 2). Linear subspace reconstruction *without* LLR regularization was used to assess the SNR and g-factor of these encodings. One instance of the g-factor for the reconstruction with LLR (𝜆𝜆=0.001) was presented to provide a sense of noise level after LLR regularization (note that the g-factor calculation for non-linear reconstruction (e.g., those using LLR regularization) is specifically valid within the tested noise level and may not be universally applicable to other scenarios due to variations in noise and image characteristics (69)). All empirical g-factor and SNR maps were calculated using the pseudo-multiple replica method (70) with 60 replicas. The ss-EPTI with *k-t* partial Fourier sampling (PF factor ∼0.85) was evaluated to improve the reconstruction conditioning at higher spatial resolutions (1.5-mm and 1.25-mm).

### 3.2 ss-EPTI image quality and tSNR evaluation

To evaluate ss-EPTI compared to ss-EPI, two ss-EPTI protocols were acquired using a gradient-echo sequence. The first protocol at a higher spatial resolution (2-mm isotropic) was acquired to test ss-EPTI’s capability to mitigate image distortion, a challenge more pronounced at higher spatial resolution in EPI. The second protocol at a lower resolution (3-mm iso) was acquired to test its effectiveness in alleviating severe signal dropout, a common issue at lower resolutions in EPI. The 2-mm protocol was acquired with the following imaging parameters: FOV = 216×216×116 mm^3^ (readout × PE × slice), number of echoes = 105, TE_range_ = 2.6–60.8 ms, ESP = 0.56 ms, TR = 2.3 s, multiband = 2, no *k-t* partial Fourier. The 3-mm protocol was acquired with: FOV = 216×216×126 mm^3^ (readout × PE × slice), number of echoes = 125, TE_range_ = 2.9–63.7 ms, ESP = 0.49 ms, TR = 1.7 s, multiband = 2, no *k-t* partial Fourier. For comparison, corresponding ss-EPI data at the same spatial resolutions and coverages were acquired with the following parameters: the 2-mm iso EPI protocol acquired with an in-plane acceleration = 2, ESP = 0.55 ms, TR/TE = 2000/35 ms, multiband = 2; and the 3-mm iso EPI protocol acquired with ESP = 0.49 ms, TR/TE = 1500/35 ms, multiband = 2. Note that the TR values of all EPI protocols were minimized to reduce their dead time and ensure fairness during the comparison of tSNR efficiency (SNR/√TR). The TR values of EPTI protocols were chosen to achieve optimized readout length for high SNR efficiency and therefore are slightly longer than the corresponding EPI protocols. The 2-mm ss-EPI and ss-EPTI protocols were used to evaluate the tSNR efficiency, and, for comparison, 3-shot EPTI data were also acquired with the same spatial parameters, a volume TR 3 times that of the single-shot acquisition, and an ESP = 0.67 ms due to the larger blip size required. Sixty dynamics were acquired for tSNR calculation. A T_2_*- based weighted optimal combination of the multi-echo images (the ‘weighted-combined’ image) was obtained by using the method described in previous study (11) for BOLD fMRI. The per-voxel T * value was estimated through a mono-exponential fit of the multi-echo ss-EPTI images.

### 3.3 Multi-echo fMRI using ss-EPTI

To demonstrate the use of ss-EPTI in fMRI, both breath-hold task and resting-state fMRI data were acquired using the 3-mm protocol described above. For the breath-hold task, ss-EPTI were acquired for a total of 5.6 minutes, during which the volunteer followed a paradigm consisting of 4 repetitions of breath-holding alternating with periods of paced breathing (8 repetitions of 3-s inspiration and 3-s expiration) as shown in Fig. 6. The breath-holding period was always followed by expiration. For the resting-state fMRI, ss-EPTI were acquired for a total of 12 minutes. Slice-timing correction and motion correction were performed using AFNI (71). Registration to the T_1_-weighted anatomical reference image (MPRAGE acquired at 1-mm iso resolution) was performed using FSL (72–74). The fMRI data were smoothed with a Gaussian kernel with 5-mm FWHM. For the breath-hold task fMRI data, activation (*t*-score) maps were obtained using a standard GLM as implemented in FSL FEAT (75). Regressors for the task blocks were modeled as a square function convolved with an HRF (76,77). For the resting-state data, ME-ICA (17,60) was performed using *tedana* (61) to demonstrate the feasibility of using ss-EPTI’s multi-echo images to extract TE-dependent information and differentiate BOLD signal from non-BOLD physiological noise.

**FIG. 6.**
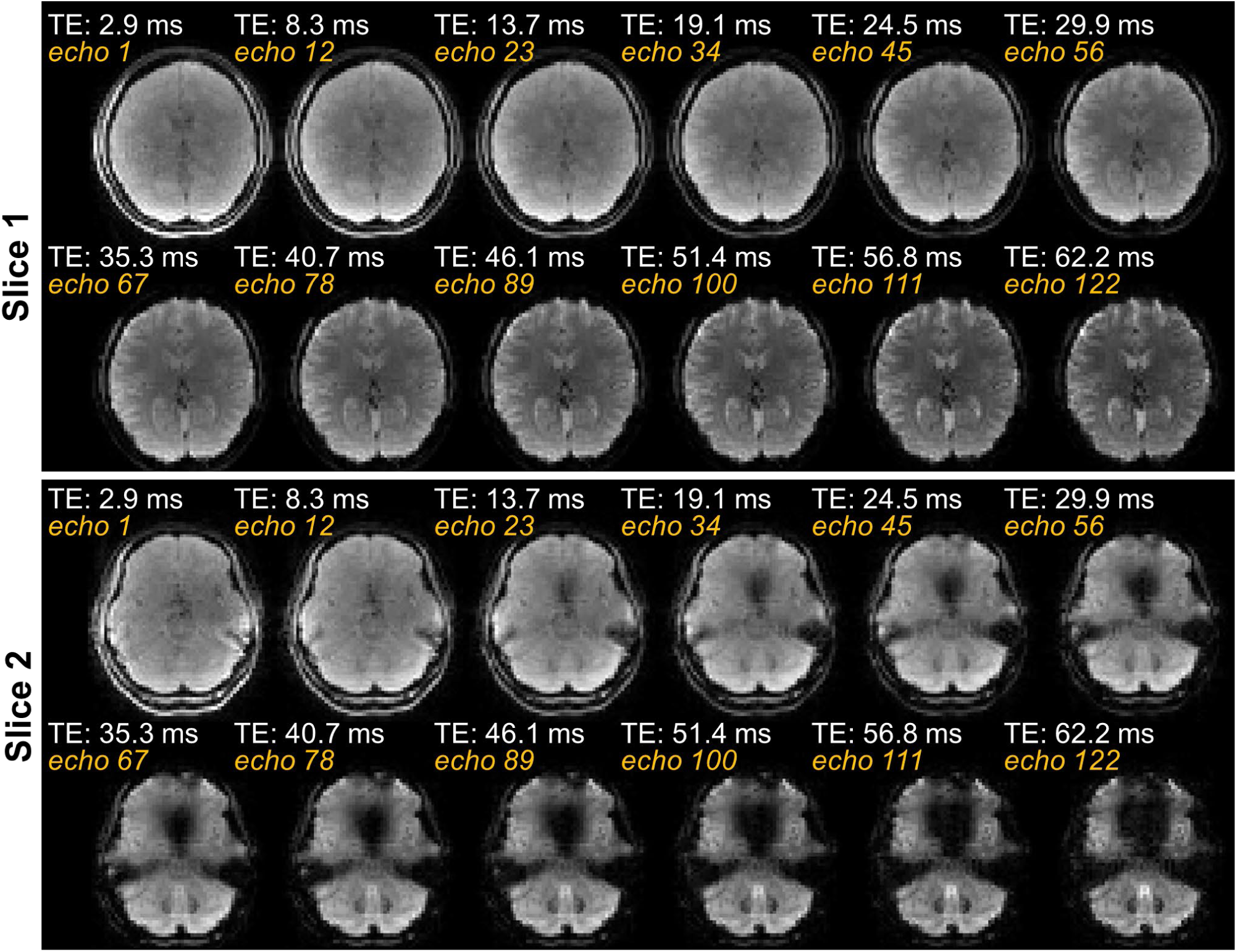
Multi-echo images (10 out of a total of 125 echoes) acquired at 3-mm isotropic resolution, with whole-brain coverage (42 slices), with TE values from 2.9 ms to 62.2 ms, obtained by ss-EPTI in a volume TR of 1.7 s. Two slices are shown, including one from the bottom part of the brain to demonstrate ss-EPTI’s performance in challenging areas with large *B*_0_ inhomogeneity. The contrast changes across echoes from PD-weighted to T_2_* weighted, and the rapid signal decay in the large *B*_0_ inhomogeneity areas, are well captured and resolved. Notably, the early echoes effectively recover the signal intensity dropout observed in later TEs.

### 3.4 Distortion-free diffusion MRI using ss-EPTI

To evaluate the data quality for dMRI, *in-vivo* whole-brain diffusion data were acquired using ss-EPTI at 1.5- mm isotropic resolution in a total scan time of 12 minutes. The imaging parameters are: *b* = 1000 s/mm^2^ and 2000 s/mm^2^, 64 directions for each b value, along with 16 *b* = 0 s/mm^2^ images (144 volumes in total); FOV = 216×216×99 mm^3^ (readout × PE × slice), number of echoes = 82, TE_range_ = 52–105 ms, ESP = 0.66 ms, TR = 5 s, multiband = 2, *k-t* partial Fourier = 0.75. The multi-echo diffusion images were averaged to i) an all-echo-combined dataset and ii) 5 different TE data (averaged 16 echoes for each data) for diffusion tensor imaging (DTI). The ss-EPTI dMRI data were processed using FSL (72–74,78,79) including FLIRT and DTIFIT. Since EPTI is free from both field-inhomogeneity-induced distortion and eddy-current distortion, no corrections were performed.

### 3.5 ss-EPTI at ultra-high field 7T

The stronger field inhomogeneity at 7T leads to severe distortion in EPI, and the shorter T_2_* makes it challenging to acquire more echo images without impractically high parallel imaging acceleration factors. To demonstrate the effectiveness of ss-EPTI for distortion-free multi-echo imaging at 7T, whole-brain gradient-echo data were acquired at 1.5-mm isotropic resolution with the following parameters: FOV = 216×216×105 mm^3^ (readout × PE × slice), number of echoes = 80, TE_range_ = 4.4–54 ms, ESP = 0.63 ms, TR = 2.5s, multiband = 2, *k-t* partial Fourier = 0.875. Conventional ss-EPI was also acquired at the same resolution with an in-plane acceleration factor of 3, TR/TE = 1900/23 ms, ESP = 0.63 ms. A whole-brain spin-echo data were also acquired at 1.25-mm isotropic resolution, with the following parameters: FOV = 210×210×78 mm^3^ (readout × PE × slice), number of echoes = 61, TE_range_ = 39–81 ms, ESP = 0.7 ms, TR = 3.1s, multiband = 2, *k-t* partial Fourier = 0.71.

## Results

The performance of single-shot EPTI encoding was characterized and optimized through simulation experiments as shown in Fig. 2. Figure 2A shows the comparison between the conventional and the proposed single-shot encoding. The g-factor, SNR, and 10× error maps of the weighted-combined images using the two encodings are shown for comparison. The proposed encoding markedly reduces the g-factor, especially in regions with large *B*_0_ offsets. It achieves a reduced ESP (0.56 ms vs. 0.93 ms at 2-mm resolution), which minimizes dead time and enables denser sampling (107 echoes vs. 65 echoes in the same 60 ms readout duration). This denser sampling, in combination with the reduced g-factor, results in a higher SNR than the conventional encoding in the single-shot regime (90.8 vs. 63.9). Lower reconstruction errors (nRMSE of 1.61% vs. 4.83%) are also observed using the new encoding.

To optimize the single-shot encoding, different spatial (*R*_PE_) and temporal (*R*_T_) acceleration factors at 3-mm, 2- mm, and 1-5 mm spatial resolutions were evaluated, and their g-factor maps are shown in Fig. 2B. As anticipated, increasing spatial resolution leads to elevated g-factor as shown in Fig. 2B because of the increased acceleration in *k-t* space due to a larger matrix size and a longer ESP. At a certain spatial resolution, both the ‘spatial-favored’ configuration (a small *R*_PE_=4 and a high *R*_T_), and the ‘temporal-favored’ configuration (a high *R*_PE_=8 and a small *R*_T_) yielded higher g-factors, while the encoding with *R*_PE_ = 6 consistently achieved the lowest g-factor for all tested resolutions, indicating a balanced spatiotemporal acceleration strategy at these resolutions. The introduction of the LLR constraint to the linear subspace reconstruction further reduced the g-factor due to improved reconstruction conditioning. The histograms of the g-factor maps corroborate these findings. Figure 2C shows that employing *k-t* partial Fourier in ss-EPTI further improved the accuracy of reconstruction at higher spatial resolution, resulting in lower nRMSEs at both 1.5-mm and 1.25-mm resolutions, with more pronounced improvements for the 1.25-mm case with a higher acceleration, as expected.

Figures 3A and 3B evaluate the image quality of ss-EPTI compared to ss-EPI acquired at comparable temporal resolutions. At 2-mm resolution, ss-EPI exhibits severe geometric distortion even with the use of a parallel imaging acceleration factor of 2, while ss-EPTI produces distortion-free images with well-maintained anatomical integrity. At 3-mm, gradient-echo ss-EPI has pronounced signal dropout in brain areas with large *B*_0_ inhomogeneity. In contrast, ss-EPTI effectively recovers the signal dropout by leveraging its early TE information in the optimal T_2_*-weighted echo combination.

**FIG. 3.**
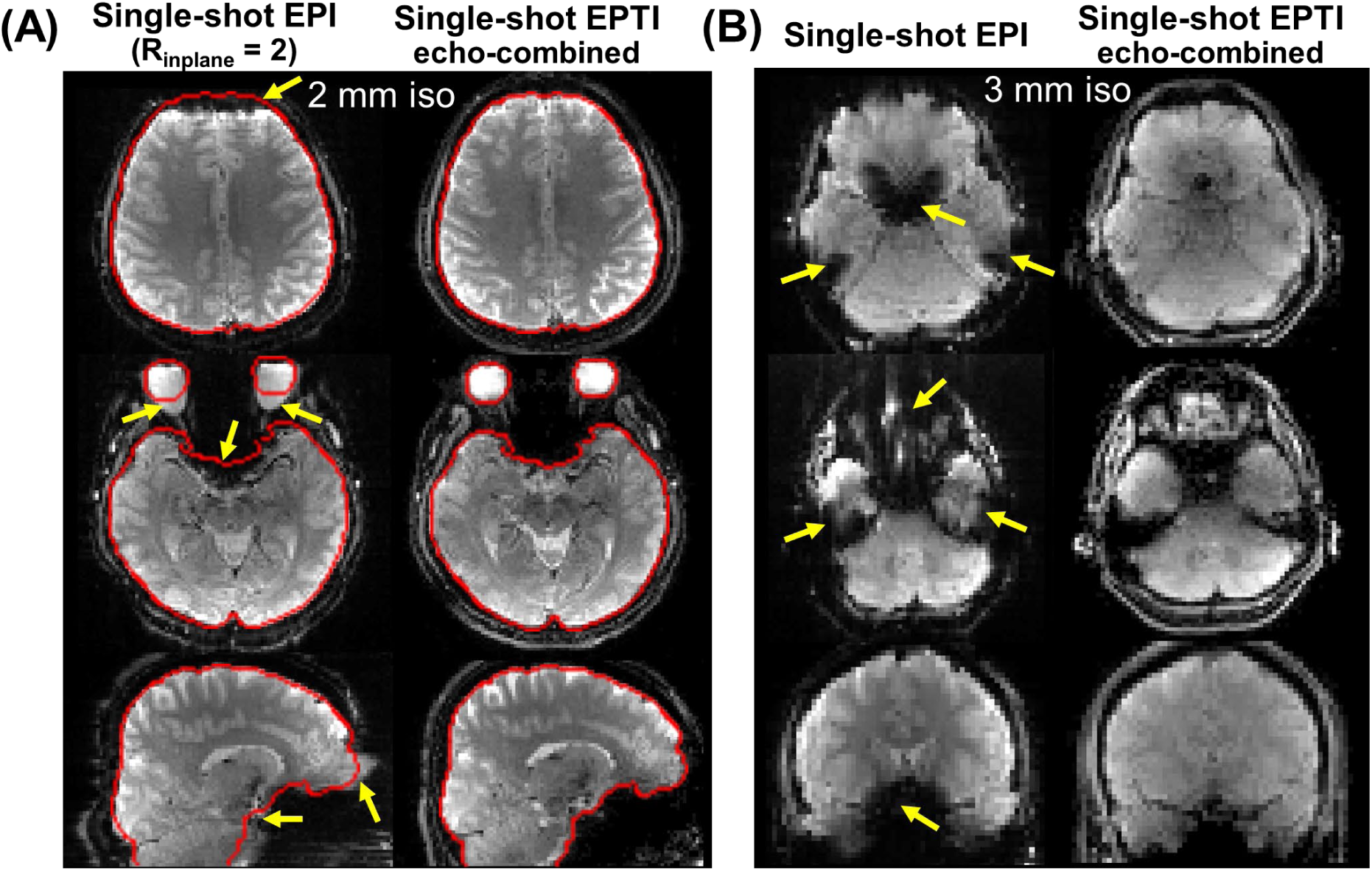
Evaluation of image quality of ss-EPI and ss-EPTI at 2-mm (**A**) and 3-mm (**B**) isotropic resolutions, acquired at comparable temporal resolutions. EPI shows severe geometric distortion even with the use of a parallel imaging acceleration factor of 2 at 2 mm, and pronounced signal dropout in brain areas affected by large *B*_0_ inhomogeneity at 3 mm, as indicated by the yellow arrows. In contrast, ss-EPTI provides distortion-free imaging with good anatomical integrity, and effectively recovers the signal dropout in challenging areas. The red contour represents the edge of the brain estimated from the EPTI data, to aid in evaluating differential geometric distortion. The presented images are optimal T_2_*-weighted echo-combined ss-EPTI images.

Figure 4 illustrates a comparison of the tSNR efficiency (tSNR/√TR) among gradient-echo ss-EPI, ms-EPTI, and ss-EPTI without and with LLR regularization. Compared to ss-EPI, ms-EPTI demonstrates significantly higher tSNR efficiency in challenging brain areas (e.g., temporal lobes, highlighted by white arrows), attributed to EPTI’s capability to acquire early TEs images with recovered signal dropout. However, ms-EPTI exhibits an overall lower tSNR efficiency, because of the inherently higher vulnerability to motion/physiological noise of multi-shot acquisition compared to single-shot, especially at low to moderate spatial resolutions and in gradient-echo sequences. Benefiting from the improved robustness against motion/physiological noise using a single-shot acquisition, ss-EPTI provides markedly higher tSNR than ms-EPTI. Compared to ss-EPI, ss-EPTI now results in higher overall tSNR efficiency in addition to the substantial improvement in challenging areas due to its more efficient readout. The use of LLR regularization further increases the SNR over linear reconstruction.

**FIG. 4.**
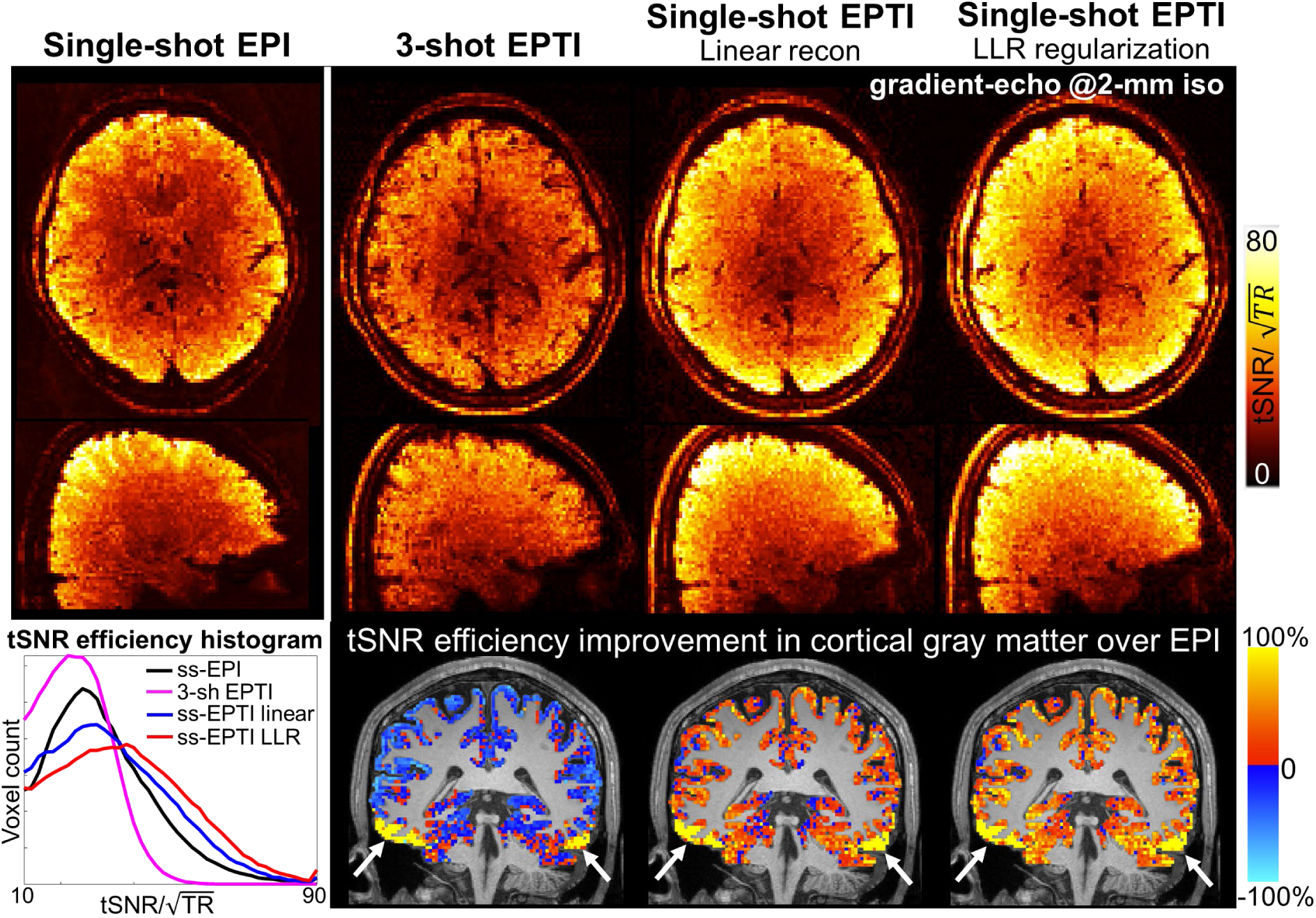
Comparison of tSNR efficiency (tSNR/√TR) among gradient-echo ss-EPI, ms-EPTI (3-shot), and ss-EPTI without and with LLR regularization at 2-mm isotropic resolution. The maps and the histograms of tSNR efficiency for all cases are depicted for comparison. tSNR efficiency improvement maps in cortical gray matter, where signals are more susceptible to physiological noise and partial volume changes, are shown, calculated by taking the ratio of the tSNR efficiency maps of EPTI and ss-EPI and subtracting one. As can be seen, ms-EPTI achieves significantly higher tSNR efficiency in challenging brain areas (highlighted by white arrows), albeit with an overall lower tSNR compared to ss-EPI due to the relatively high vulnerability to physiological noise of multi-shot acquisition. ss-EPTI achieves improved tSNR efficiency over ms-EPTI due to improved motion/physiological robustness. Compared to ss-EPI, ss-EPTI now achieves a higher overall tSNR efficiency due to the improved readout efficiency, in addition to the substantial improvement in challenging brain areas attributed to the early TE information.

The ability of ss-EPTI to provide time-series data free from distortions can further improve temporal stability by mitigating dynamic distortions commonly observed in EPI. The dynamic distortions of EPI are caused by field changes due to drift, motion, or physiological dynamics. They can often resemble motion (image shifts along the PE direction) and lead to signal variations. Figure 5 presents a comparison between ss-EPI and ss-EPTI during acquisitions with large field changes induced by periods of paced breathing in a 5.6-minute breath-holding fMRI task. For each subject, the estimated rigid-motion parameters from AFNI, and the carpet plots of the signal normalized by the mean signal of each voxel, are plotted. The motion parameters of ss-EPI show ‘apparent motion’ during paced breathing (image shifts along the PE direction, plotted in red) due to breathing-induced field changes. In comparison, ss-EPTI demonstrates minimal fluctuations due to this effect. Residual image shifts observed in subject 1 are likely attributed to actual rigid head movements accompanying paced breathing, and a similar pattern is also present in other motion parameters. In addition, as can be seen in the carpet plots, the signal intensity variations caused by these dynamic distortions in EPI were effectively mitigated in ss-EPTI. Consistent conclusions can be drawn across three subjects exhibiting varying levels of motion.

**FIG. 5.**
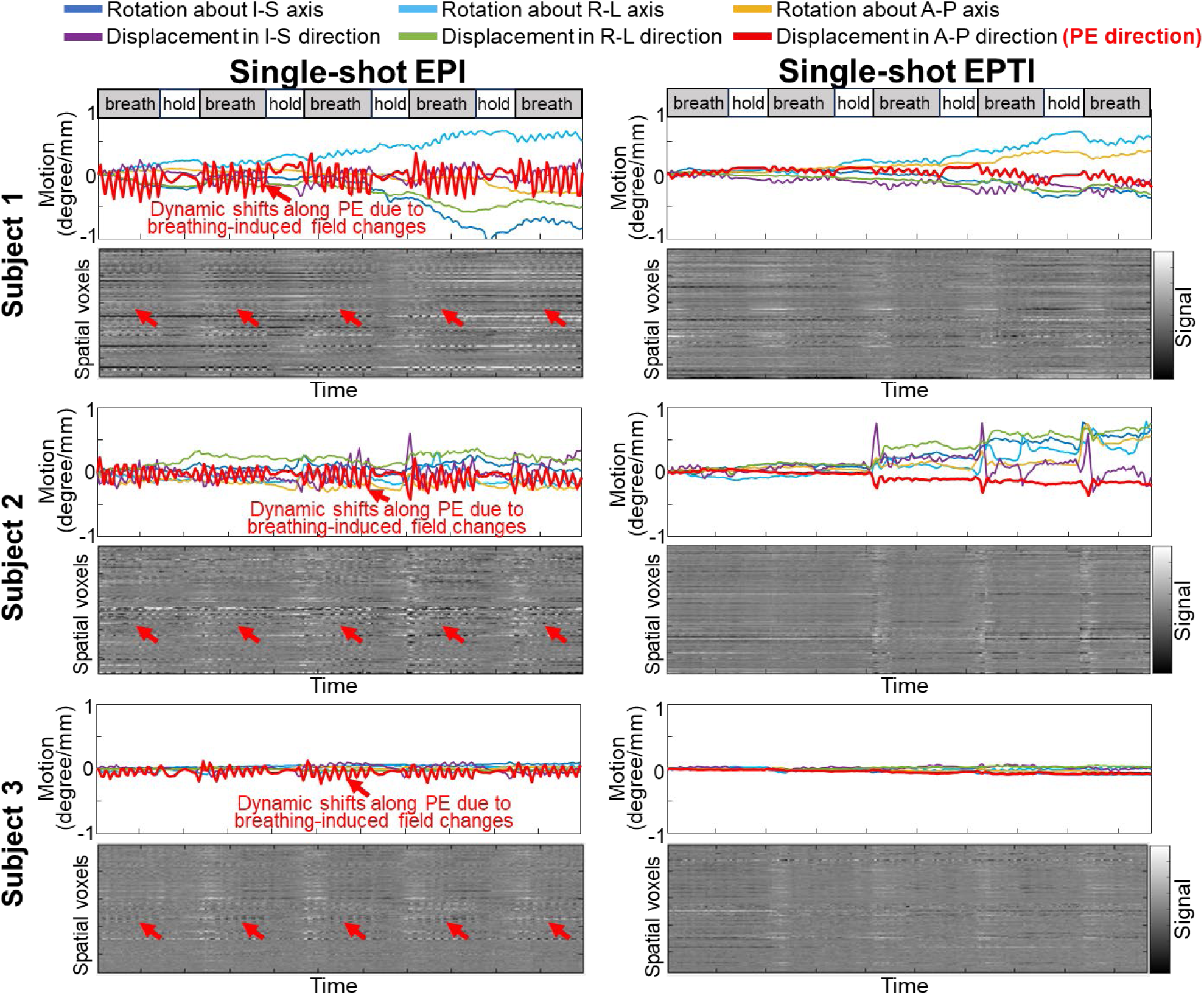
Evaluation of the ability of ss-EPTI to mitigate dynamic distortions and associated signal variations, by comparing it with ss-EPI during acquisition runs with *B*_0_ field changes induced by periods of paced breathing in a 5.6-min breath-holding fMRI task. The timing paradigm for paced breathing and breath-holding is shown at the top of the figure. The 6 estimated rigid-motion parameters from AFNI and the corresponding carpet plots of the image data across time, normalized by the mean signal of each voxel, are plotted for three subjects. While ss-EPI shows ‘apparent motion’ (image shifts along the phase-encoding direction, plotted in red) during paced breathing due to breathing-induced field changes, ss-EPTI shows minimal fluctuations due to this effect. Any residual image shifts observed in subject 1 are likely attributed to actual rigid head movements accompanying paced breathing. As shown in the carpet plot, the variations of the signal intensity caused by these dynamic distortions in EPI were effectively mitigated in ss-EPTI.

Figure 6 shows the 10 selected multi-echo images out of the total of 125 echoes resolved across the ss-EPTI readout, which was acquired at 3-mm isotropic resolution with whole-brain coverage (42 slices) in a volume TR of 1.7 s. Starting from a very early TE of 2.9 ms and extending to a TE of ∼60 ms, the image contrasts changing from proton-density-weighted to T *-weighted are well captured. In the bottom slice where large *B*_0_ inhomogeneity exists, rapid signal decay is resolved even in challenging areas with signal dropout, indicating a good representation of signal evolution by the reconstruction. The early TEs effectively recover the severe dropout in signal intensity observed in later TEs.

Figures 7 and 8 demonstrate the use of ss-EPTI for multi-echo fMRI for task and resting-state fMRI. Figure 7 shows the functional sensitivity of different ss-EPTI echoes in a breath-holding task experiment from 3 subjects. The multi-echo ss-EPTI data show that the functional sensitivity changes across different echoes, and the changes are consistent across all subjects. Specifically, the early echoes demonstrate an overall lower *t*-score compared to the later echoes as expected, but yield significantly higher *t*-scores in challenging brain regions (white arrows and zoom-in views) whose functional sensitivity can be irrevocably lost at a typical BOLD TE of ∼30–40 ms. These brain regions align well with the short-T * areas identified in the T * maps obtained by ss-EPTI. This demonstrates the ability of the multi-echo images acquired by ss-EPTI to not only obtain good signal intensity and signal evolution information, but also capture the subtle and different functional signal changes across TEs.

**FIG. 7.**
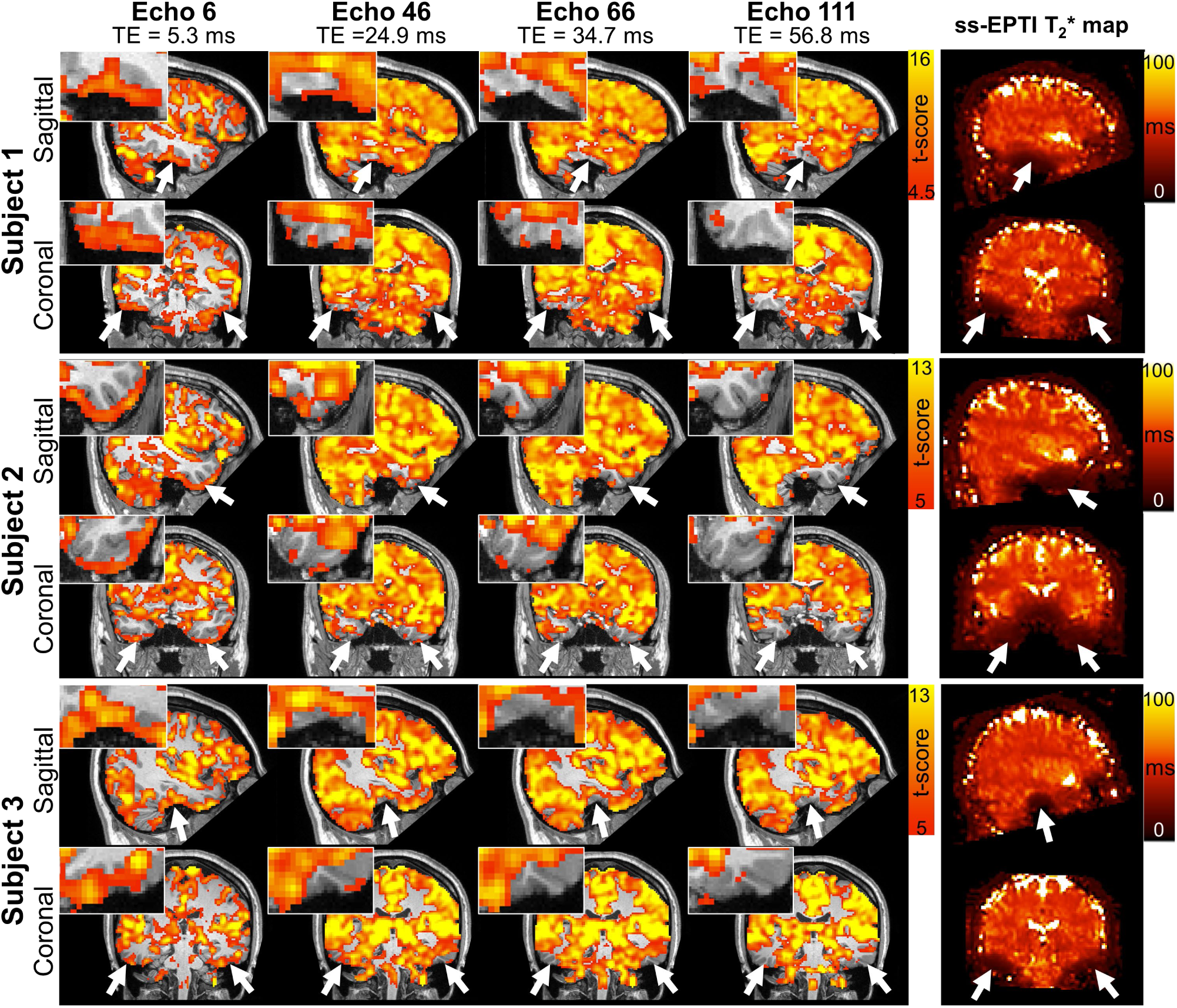
Results of the multi-echo fMRI experiment acquired by ss-EPTI at 3-mm isotropic resolution during a breath-hold task from 3 subjects. T-score activation maps of 4 different echoes are shown in both sagittal and coronal views. Additionally, T_2_* maps fitted from the ss-EPTI multi-echo images are shown. The multi-echo images of ss-EPTI show expected functional sensitivity changes across different echoes, consistently observed in all subjects: the early echoes provide an overall lower *t*-score but yield significantly higher t-scores in challenging temporal lobe areas than the later TEs (e.g., typical BOLD TE of 30–40 ms). These areas, highlighted in white arrows with zoomed-in views, align well with the short T_2_* areas identified in the T_2_* maps. This indicates the ability of the multi-echo images acquired by ss-EPTI to not only obtain good signal evolution across TEs, but also capture the subtle functional signal changes across TEs.

**FIG. 8.**
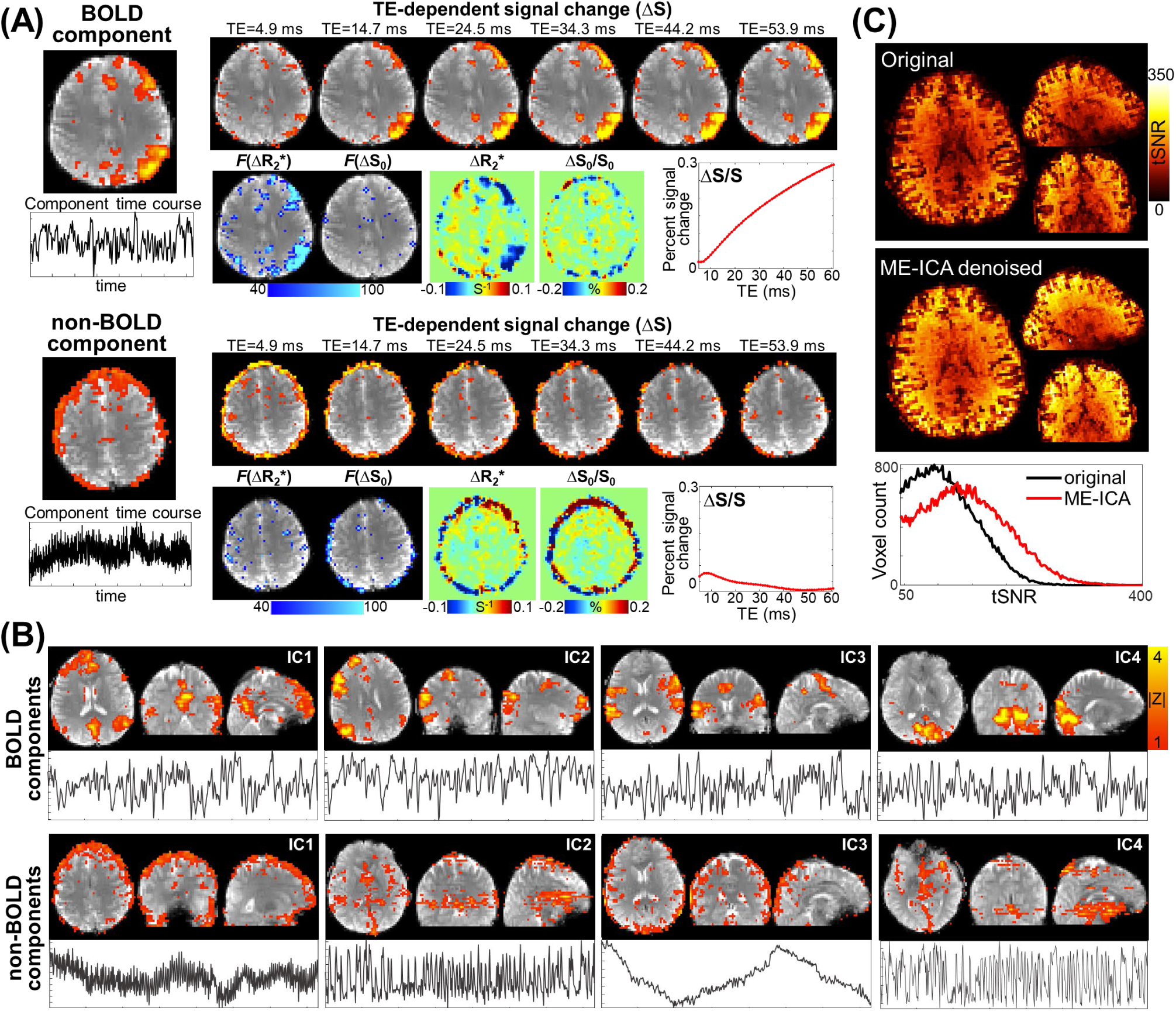
Results of the resting-state multi-echo fMRI experiment acquired by ss-EPTI at 3-mm isotropic resolution, where ME-ICA was applied to the ss-EPTI multi-echo data. (**A**) Spatial and temporal patterns (left), and TE-dependency (right) of an example BOLD (top) and non-BOLD (bottom) component extracted from ss-EPTI data. The TE-dependency of each component is illustrated through signal change across different TEs (top), the percent signal change as a function of TE (bottom right plot), the goodness of fit of the multi-TE signal to the R_2_* and S_0_ signal models (bottom left, *F*(ΔR_2_*) and *F*((ΔS_0_) maps), and the quantitative R_2_* and S_0_ changes (bottom middle, ΔR_2_* and ΔS_0_/S_0_ maps). The BOLD component exhibits a linear relationship between percent signal change (ΔS/S) and TE, a strong fit to the R_2_* signal model (*F*(ΔR_2_*)), and localization corresponding to large quantitative R_2_* changes (ΔR_2_*). In contrast, the non-BOLD component display relatively constant percent signal change (ΔS/S) across TEs (signal change ΔS decays exponentially), a strong fit to the S_0_ signal model (*F*(ΔS_0_)), and localization corresponding to large quantitative S_0_ changes (ΔS_0_/S_0_). (**B**) Examples of additional BOLD and non-BOLD components successfully distinguished using ss-EPTI data. BOLD components exhibit network-like spatial patterns and relatively lower-frequency time courses, while non-BOLD components have artifacts-like spatial patterns and higher-frequency time courses. (**C**) tSNR maps of ss-EPTI data (optimal T_2_*-weighted echo-combined image) before and after ME-ICA denoising, along with their corresponding tSNR histogram plots. Improved tSNR is observed after removing non-BOLD physiological noise components using ss-EPTI combined with ME-ICA.

Figure 8 assesses the ability of ss-EPTI’s multi-echo images to extract TE-dependent information about signal fluctuations in a resting-state fMRI experiment. By applying ME-ICA, BOLD signal components were successfully distinguished from non-BOLD signals in the ss-EPTI data, with distinct spatial and temporal patterns and TE-dependency (Fig. 8A & 8B). BOLD components have relatively lower-frequency time courses and network-like spatial patterns. They exhibit a linear relationship between percent signal change (ΔS/S) and TE, and a strong goodness of fit to the R_2_* signal model (*F*(ΔR_2_*)). The localization of the signal change also corresponds to large quantitative R * changes (ΔR *). In contrast, non-BOLD components have higher-frequency time courses and artifacts-like spatial patterns (e.g., at the brain edge or within CSF). They exhibit relatively constant percent signal change (ΔS/S) across TEs (signal change ΔS decays exponentially), and a strong goodness of fit to the S_0_ signal model (*F*(ΔS_0_)). The localization of the signal changes also corresponds to large quantitative S_0_ changes (ΔS_0_/S_0_). By identifying and removing the non-BOLD components, unwanted time-series variance was further reduced, resulting in improved tSNR for resting-state fMRI (Fig. 8C).

Figure 9 and 10 present the diffusion MRI data acquired by ss-EPTI at 1.5-mm isotropic resolution with whole-brain coverage (66 slices), where 128 diffusion encodings and 16 b=0 s/mm^2^ images were acquired in 12 minutes. The single-direction diffusion-weighted images and mean DWIs (Fig. 9) at *b*=1000 s/mm^2^ and *b*=2000 s/mm^2^ show high image quality and high SNR without any noticeable geometric distortion or ghosting artifacts. The DWIs across all directions and shells exhibit identical geometry aligning with the same red contour extracted from mean DWI, demonstrating the elimination of the dynamic distortion caused by diffusion encoding dependent eddy currents. The high-quality FA maps (Fig. 10A) show detailed fiber structures even in the deep brain regions that typically suffer from low SNR and image distortion in EPI (zoom-in views). Multi-TE images were obtained from the same ss-EPTI dataset without extra scan time, and multi-TE diffusion-weighted images, FA, and MD maps are shown in Fig. 10B.

**FIG. 9.**
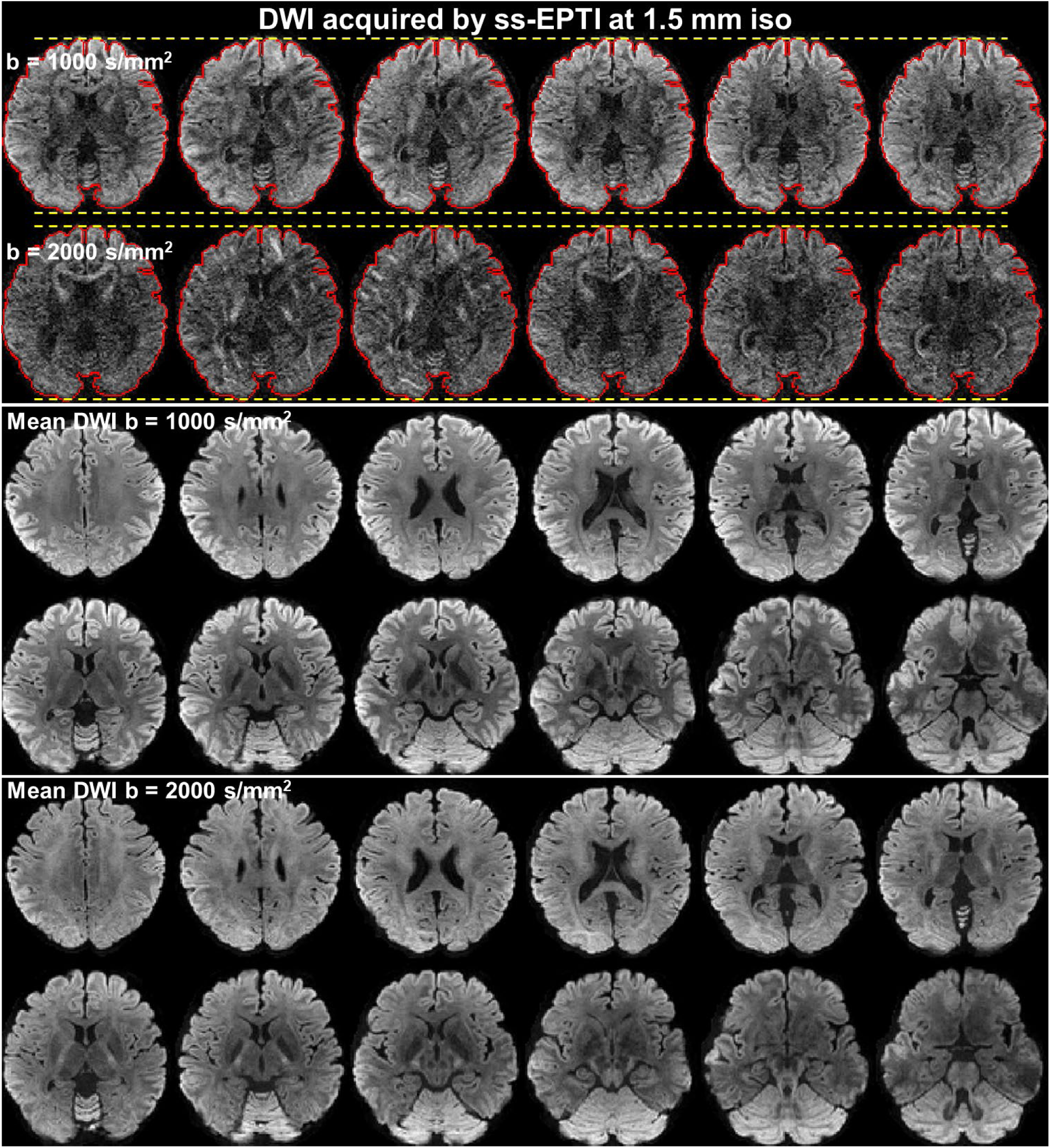
Diffusion MRI data acquired using single-shot EPTI at 1.5-mm isotropic resolution with whole-brain coverage (66 slices), where 128 diffusion encodings and 16 b=0 s/mm^2^ images were acquired in 12 minutes. Single-direction diffusion-weighted images at b=1000 s/mm^2^ and b=2000 s/mm^2^ of 6 example diffusion directions are shown (top) with an identical red contour extracted from mean DWI. The mean DWIs of the two b-values across different slices are shown in the bottom. The DWIs and mean DWIs show high image quality and high SNR without noticeable geometric distortion or artifacts. The DWIs across all directions and shells exhibit identical geometry aligning with the same red contour, demonstrating the removal of the dynamic distortion caused by diffusion encoding dependent eddy currents.

**FIG. 10.**
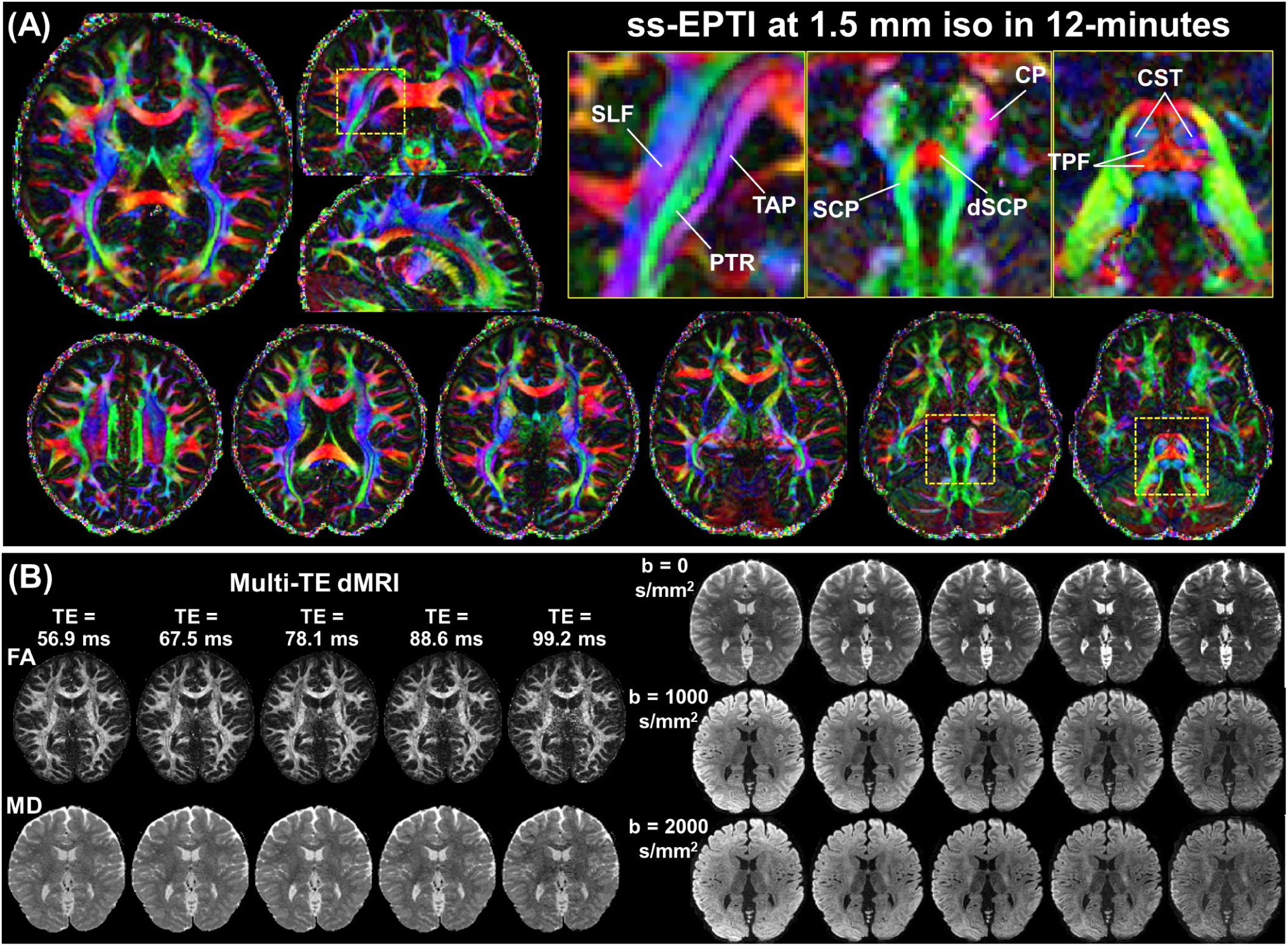
(**A**) Color-coded FA maps calculated from 12 minutes of single-shot EPTI data at 1.5 mm isotropic spatial resolution. The zoomed-in areas highlight the high SNR and fidelity of the detailed fiber structures observed in the deep brain regions that typically suffer from low SNR and image distortion in EPI. (SLF: superior longitudinal fasciculus, TAP: tapetum of the corpus callosum, PTR: posterior thalamic radiation, CP: cerebral peduncle, SCP: superior cerebellar peduncles, dSCP: decussation of the superior cerebellar peduncles, CST: corticospinal tract, TPF: transverse pontine fibers). (**B**) Multi-TE images were acquired within the same ss-EPTI dataset without extra scan time, providing high-quality multi-TE estimates of both FA and MD (mean diffusivity), as well as multi-TE DWIs.

Figure 11 examines the performance of ss-EPTI at ultra-high-field strength at 7T. Figure 11A shows gradient-echo ss-EPI and ss-EPTI at 1.5-mm isotropic resolution. While the single-shot EPI exhibits distortions even with a 3× in-plane acceleration and ghosting artifacts, ss-EPTI provides distortion-free imaging with good quality and additional multi-echo information. A spin-echo ss-EPTI acquired at 1.25-mm isotropic resolution was also shown, showing the capability of ss-EPTI for imaging at relatively higher spatial resolution.

**FIG. 11.**
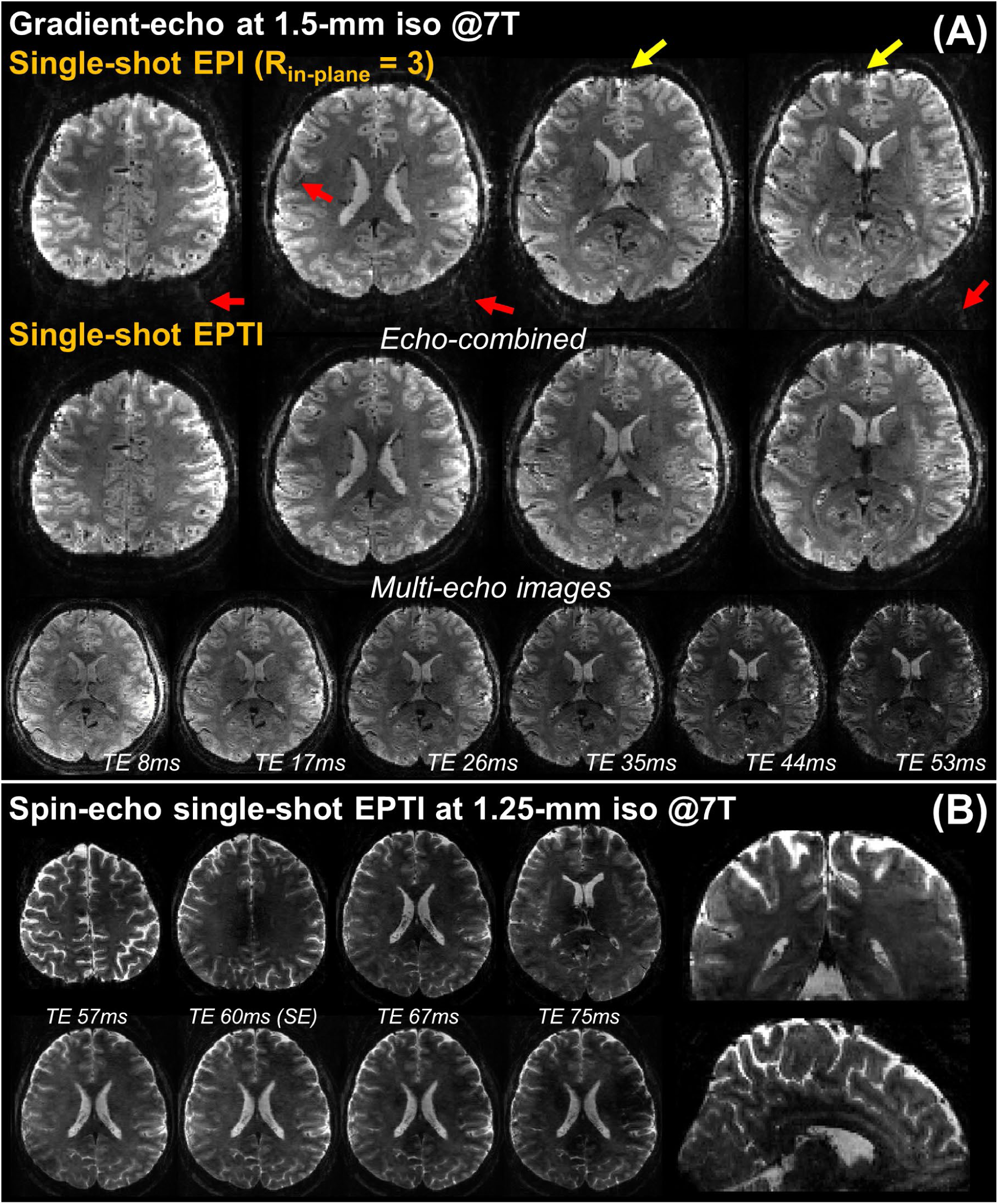
ss-EPTI dataset acquired at ultra-high-field strength (7T). (**A**) Example gradient-echo data at 1.5-mm isotropic resolution acquired using ss-EPI and ss-EPTI. ss-EPI shows severe distortions (yellow arrows) even after using a 3× in-plane acceleration with some ghosting artifacts (red arrows), while ss-EPTI provides distortion-free images with additional multi-echo information. (**B**) Example spin-echo ss-EPTI data acquired at 1.25-mm isotroipic resolution with a TR of 3.1 s.

### Discussion and Conclusion

This study develops EPTI from the original multi-shot acquisition to a single-shot scheme to achieve high robustness to motion and physiological noise, high temporal resolution, and improved tSNR efficiency. This development was motivated by the goal of broadening the applicability of EPTI in both clinical and neuroscientific contexts where these improvements are of great importance. We have demonstrated that the proposed ss-EPTI achieves improved robustness to motion/physiological noise and therefore higher tSNR efficiency than ms-EPTI at standard spatial resolutions, while inheriting EPTI’s key features of distortion-free imaging, densely-sampled multi-echo imaging, and SNR-efficient continuous readout. As a result, EPTI now provides improved image quality, richer multi-echo information, and higher acquisition efficiency—with comparable temporal resolutions compared to single-shot EPI. This offers a new acquisition tool for a variety of dynamic imaging applications where ss-EPI currently serves as the standard sequence, such as fMRI and dMRI.

Single-shot EPTI was achieved through the design of a new encoding pattern and the optimization of encodings for enhanced spatiotemporal correlation in the *k-t* space. Since only one shot is acquired to cover the *k-t* space, a stronger spatiotemporal correlation is critical to recover the highly undersampled *k-t* space. Single-shot EPTI achieved this by employing an encoding pattern that requires small gradient blips along *k_y_* and achieves minimized ESP, which effectively improves the spatiotemporal correction and reconstruction conditioning. The shorter ESP also reduces dead time and enables denser sampling within the EPTI readout, which, together with the reduced g-factor, leads to improved SNR efficiency. The reduced gradient blips also avoid large eddy currents in comparison to the conventional encoding pattern where a large gradient blip is needed to traverse through the entire *k*-space when moving to the single-shot scheme.

The interdependent spatial and temporal acceleration interact to determine the spatiotemporal correlation of the single-shot *k-t* data, and they were optimized to reach a balance for improved reconstruction conditioning. The optimized spatial acceleration factor *R*_PE_ = 6 is higher than the typical acceleration factor used in conventional parallel imaging, but still results in low g-factor and reconstruction error because the complementary sampling across different TEs aids in the recovery of *k-t* undersampling by exploiting the strong temporal correlation in addition to the spatial information provided by the coil array sensitivities. The synergistic role of multi-coil spatial information and temporal correlation in the reconstruction also explains why either sacrificing temporal correlation to reduce spatial undersampling (*R*_PE_ = 4) or sacrificing spatial correlation to reduce temporal undersampling (*R*_PE_ = 8) results in a higher g-factor than the balanced selection. Although our simulations across a range of spatial resolutions consistently yielded the same optimal parameter, we note that this optimized parameter may not encompass all possible scenarios, particularly at other spatial resolutions or field strengths. The optimization process presented in this study can serve as an example of the general concept of spatiotemporal encoding optimization for EPTI, to guide the parameter optimization in other scenarios. The *k-t* partial Fourier acquisition further enhances the spatiotemporal correlation by reducing the temporal acceleration, improving the reconstruction accuracy at higher spatial resolution (e.g., higher than 1.5 mm). The use of the LLR constraint also improves the reconstruction conditioning and reduces noise. It is worth noting that LLR regularization is applied across the different coefficient maps as a low-rank constraint across echoes, assuming the signal evolution across echoes is slow and temporally smooth/low-rank (e.g., T_2_* decay). Consequently, different dynamics over time (i.e., TR volumes in fMRI, directions in dMRI) are still independent in EPTI.

As an important application of ss-EPTI, we demonstrated its utility in fMRI. The high readout efficiency in ss-EPTI contributes to the enhanced tSNR efficiency (Fig. 4), and its distortion-free nature further improves the temporal stability by eliminating dynamic distortions commonly exist in EPI caused by field changes due to motion or physiological dynamics (e.g., breathing in Fig. 5). Moreover, ss-EPTI provides multi-echo images that capture the signal evolution with high fidelity, which can be useful for fMRI as demonstrated in the two example applications in task and resting-state fMRI. First, different TE images provide different functional contrasts, and early TEs can be used to effectively recover the signal intensity and functional sensitivity dropout in challenging brain regions. Here, breath-hold fMRI task was selected to demonstrate this as it can achieve whole-brain activation in gray matter, and allow for investigation of brain areas with different T_2_* values. Second, in resting-state fMRI, the multiple TEs provide useful information about TE dependency of functional signal fluctuations, and can be used to differentiate BOLD signal and physiological noises to further improve tSNR. While conventional fMRI data processing/analysis methods were used to demonstrate the information content provided by ss-EPTI, advanced or new data processing/analysis methods (80) tailored to EPTI’s signal and noise characteristics might be able to better exploit the wealth of information obtained from ss-EPTI acquisition. Notably, the application of deep-learning based analysis methods (81,82) holds great promise for effectively analyzing and extracting the rich information embedded within EPTI’s multi-echo/multi-contrast data, representing an exciting avenue for future exploration in maximizing the utility of ss-EPTI in fMRI.

We have also demonstrated the application of ss-EPTI in dMRI. The distortion-free feature of ss-EPTI not only preserves geometric accuracy in each DWI, but can also—similar to eliminating the dynamic distortions from breathing-induced field changes—address the dynamic distortions across different DWIs due to diffusion-encoding-dependent eddy-current-induced field changes, a common problem in dMRI using EPI. As a result, no distortion or eddy-current correction was needed for the ss-EPTI dMRI data. The dMRI images are presented with high SNR (Figs. 9 & 10) benefiting from the minimized TE and high readout efficiency. The shortened TE can also provide additional SNR gain for ultra-high-field dMRI (e.g., 7T) where T_2_ in brain tissue is inherently shorter than at conventional field strengths. Additionally, the resolved multi-echo images can also enable the acquisition of diffusion metrics across different TEs (Fig. 10) that can be useful in diffusion relaxometry analyses. Compared to multi-shot, the high robustness of ss-EPTI to motion/physiological noise ensures more practical acquisitions and robust performance, particularly in applications using high or ultra-high b-values, where the large shot-to-shot phase variations caused by the strong diffusion gradient could lead to image artifacts and data corruption without proper correction when using multi-shot acquisitions, while single-shot EPTI avoids this problem.

In this study, although single-shot EPTI was tested across a range of spatial resolution (e.g., 1.25 mm to 3 mm), further increasing spatial resolution (e.g., submillimeter resolution) will lead to higher undersampling in *k-t* space and may result in amplified noise or artifacts in the reconstructed images. Improved *B*_0_ shimming (83–85) and hardware advancements such as high-performance gradient coils (e.g., head-only gradient coils (86–88)), and/or advanced reconstruction such as deep-learning-based approaches (89–91), can further improve reconstruction conditioning and achieve higher spatial resolution using ss-EPTI. Moreover, the temporal resolution of ss-EPTI can also be further improved. The long readout length used in this study was selected to achieve optimized SNR efficiency, and a shorter TR can be simply achieved by reducing the readout length. In addition, we have successfully incorporated SMS into ss-EPTI encoding and achieved good image quality at MB=2. Further increasing the MB factor is feasible but depends on the capabilities of i) the coil sensitivity along both PE and slice directions, and ii) the temporal correlation within the readout, to reconstruct the fully-sampled *k-t* space. The compatibility of EPTI with SMS is analogous to that of parallel imaging with SMS (92), with the difference that ss-EPTI also takes advantage of the temporal correlation across echoes. As with spatial resolution, advancements in hardware and reconstruction will play a key role in achieving higher temporal resolution for ss-EPTI. Finally, non-Cartesian trajectories can also be employed to achieve single-shot time-resolved imaging (93).

Finally, while we have demonstrated the use of ss-EPTI for multi-echo fMRI and diffusion MRI, it can be readily used as a readout technique in other applications such as perfusion MRI, including techniques based on ASL (Arterial Spin Labeling), DSC (Dynamic Susceptibility Contrast) or DCE (Dynamic Contrast Enhancement) to address EPI’s limitations and obtain additional multi-echo/multi-contrast information. As one example, we obtained whole-brain distortion-free multi-contrast images in a gradient-echo-and-spin-echo sequence at 2-mm isotropic resolution and a temporal resolution of 4.4 s, obtaining dynamic T_2_* and T_2_ maps (Supporting Information Figure S1), which can be useful in perfusion imaging.

## Acknowledgements

We thank Mr. Kyle Droppa, Ms. Sarah Richter, and Ms. Estee Perelgut for their help with subject recruitment and MRI scanning support, and Drs. Jingyuan Chen and Kawin Setsompop for helpful discussion. This work was supported by the NIH NINDS *BRAIN Initiative* (U24-NS129893), NIH NIA (K99-AG083056), and NIBIB (P41-EB030006 and R01-EB019437), and by the MGH/HST Athinoula A. Martinos Center for Biomedical Imaging, and was made possible by the resources provided by NIH Shared Instrumentation Grant S10-OD023637.

## Supporting Information

**GESE ss-EPTI for dynamic T_2_ and T_2_* mapping:** The efficient sampling of multi-contrast images using EPTI facilitates the rapid acquisition of quantitative maps, as demonstrated in previous studies. The transition to a single-shot acquisition significantly enhances the temporal resolution of EPTI for dynamic quantitative mapping. As a proof-of-concept experiment, multi-contrast gradient-echo-and-spin-echo (GESE) images were acquired using a GESE single-shot EPTI sequence at 2-mm isotropic resolution to achieve fast T_2_ and T * mapping with a volume TR of 4.4 s. The other imaging parameters are: FOV = 216 × 216 × 100 mm^3^ (readout × PE × slice), number of echoes = 221 (80 GEs, 141 SEs), TE_range_ of GE = 3–47 ms, TE_range_ of SE = 79–157 ms, ESP = 0.56 ms, TR = 4.4 s, multiband = 2; no partial Fourier.

**FIG. S1.**
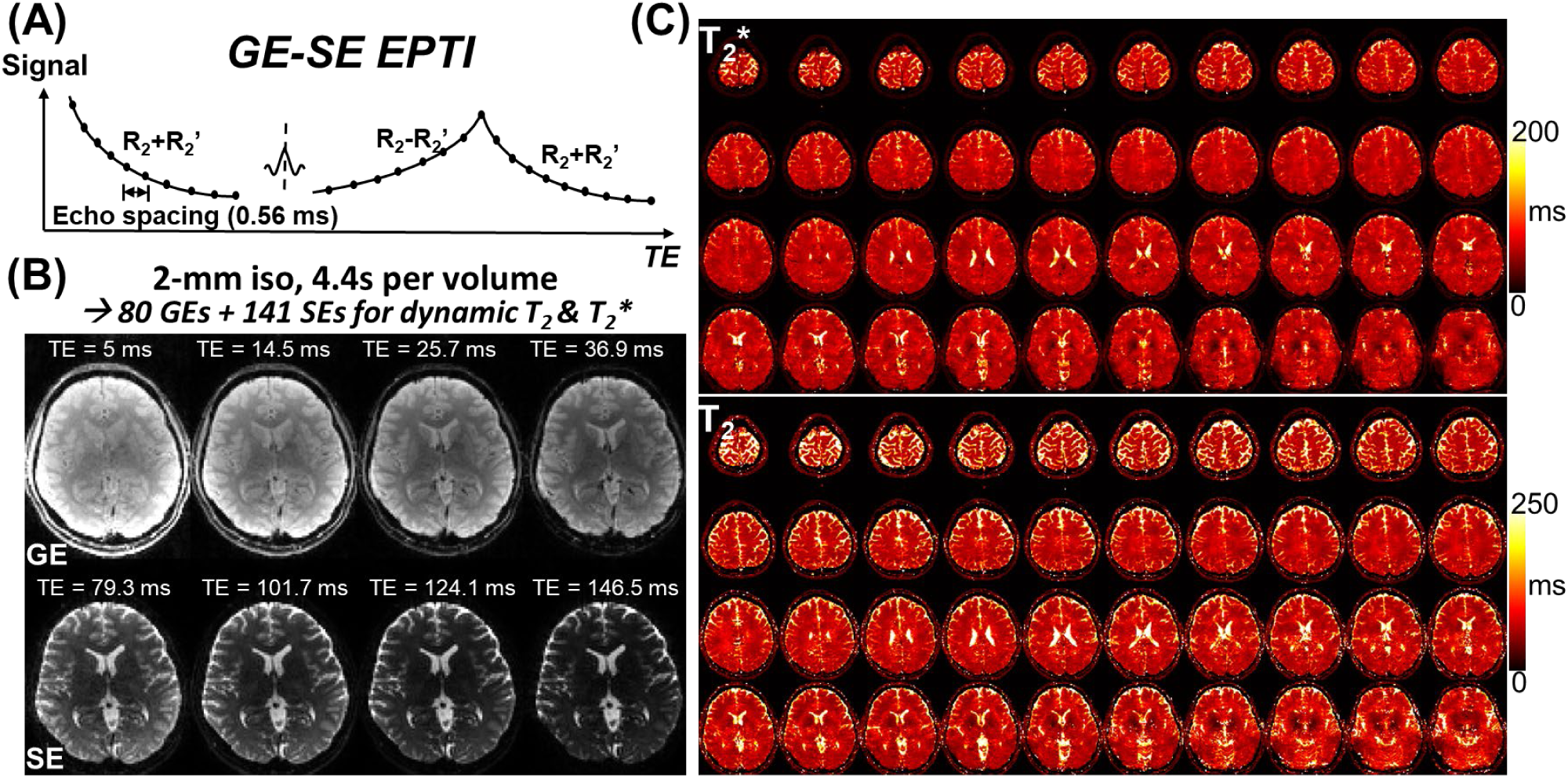
Gradient-echo-and-spin-echo (GESE) single-shot EPTI results. (**A**) Illustration of the signal evolution of a GESE ss-EPTI sequence. (**B**) Representative multi-contrast images acquired by GESE ss-EPTI at 2-mm isotropic resolution with whole-brain coverage, where 80 GE volumes and 141 SE volumes were acquired within a volume TR of 4.4s. The first row shows 4 example GE images out of the 80 GEs, and the second row shows 4 example SE images out of the 141 acquired. (**C**) Fitted T_2_* and T_2_ maps from a single dynamic across different slices are shown.

## Notes

### Competing Interest Statement

The authors have declared no competing interest.

